# Circulating BMP9 protects the pulmonary endothelium during inflammation-induced lung injury in mice

**DOI:** 10.1101/2020.05.12.088880

**Authors:** Wei Li, Lu Long, Xudong Yang, Zhen Tong, Mark Southwood, Paola Caruso, Paul D Upton, Peiran Yang, Geoffrey A Bocobo, Ivana Nikolic, Angelica Higuera, Richard M Salmon, He Jiang, Katharine M Lodge, Kim Hoenderdos, Rebecca M Baron, Paul B Yu, Alison M Condliffe, Charlotte Summers, Edwin R Chilvers, Nicholas W Morrell

## Abstract

**Rationale:** Pulmonary endothelial permeability contributes to the high-permeability pulmonary edema that characterizes acute respiratory distress syndrome (ARDS), which carries a high mortality. Circulating bone morphogeneic protein 9 (BMP9) is emerging as an important regulator of pulmonary vascular homeostasis.

**Objective:** To determine whether endogenous BMP9 plays a role in preserving pulmonary endothelial integrity, and whether loss of endogenous BMP9 occurs during lipopolysacharride (LPS)-induced lung inflammation and permeability.

**Methods:** A BMP9-neutralizing antibody was administrated to healthy adult mice and lung vasculature was examined. Potential mechanisms were delineated by transcript analysis in human primary lung endothelial cells. Impact of BMP9 was evaluated in a murine acute lung injury model induced by inhaled LPS. Levels of BMP9 were measured in plasma from patients with sepsis and endotoxemic mice.

**Main Results:** Subacute neutralization of endogenous BMP9 in mice resulted in increased lung vascular permeability, interstitial edema and neutrophil extravasation. In lung endothelial cells, BMP9 regulated a programme of gene expression and pathways controlling vascular permeability and cell membrane integrity. Augmentation of BMP9 signalling in mice with exogenous BMP9 prevented inhaled LPS-caused lung injury and edema. In endotoxemic mice, endogenous BMP9 levels were markedly reduced, due to a transient reduction in hepatic BMP9 mRNA expression and increased elastase activity in plasma. In human sepsis patients, circulating levels of BMP9 were also markedly reduced.

**Conclusions:** Endogenous circulating BMP9 is a pulmonary endothelial protective factor, down-regulated during inflammation. Supplementation with exogenous BMP9 offers a potential therapy to prevent increased pulmonary endothelial permeability in the setting of lung injury.

**Short summary:** *Scientific Knowledge on the Subject:* Increased pulmonary endothelial permeability is a major factor in the development of acute respiratory distress syndrome (ARDS). Evidence is emerging that circulating BMP9, secreted from the liver, might protect the pulmonary endothelium from injury. For example, loss of BMP9 levels or signalling receptor contributes to the development of pulmonary arterial hypertension. The role of endogenous BMP9 in endothelial permeability remains unclear.

*What This Study Adds to the Field:* Here we show that subacute neutralization of endogenous BMP9 leads to lung vascular injury, including enhanced permeability and neutrophil extravasation. BMP9 levels were markedly reduced in the setting of inflammation in mice and humans. Conversely, exogenous supplementation of BMP9 protected the lung from LPS-induced injury. This study suggests that exogenous BMP9 could offer a novel approach to prevent increased pulmonary endothelial permeability in the setting of lung injury and ARDS.

## INTRODUCTION

Endothelial dysfunction, inflammation and increased capillary permeability play central roles in the pathobiology of sepsis and the acute respiratory distress syndrome (ARDS)(1). Previous studies have identified important signaling pathways and protein-protein interactions within inter-endothelial junctions that regulate endothelial barrier function(2). However, such knowledge has not yet resulted in approved drugs that target the increased vascular permeability present in sepsis and ARDS(1), conditions associated with an unacceptably high mortality. Exploring new pathways that preserve endothelial integrity may hasten the discovery of novel approaches to the treatment of these conditions.

Bone morphogenetic protein 9 (BMP9) is a member of the transforming growth factor β (TGFβ) family that signals selectively in endothelial cells via a receptor complex comprising the high affinity type 1 BMP receptor activin receptor-like kinase 1 (ALK1), and the type 2 BMP receptors, BMP receptor type II (BMPRII) or activin receptor type 2A(3-5). ALK1 is expressed almost exclusively on endothelial cells(6), and its expression is 10 to 200-fold higher in lung compared to other tissues, indicating a particular role for ALK1-mediated signaling in homeostasis of the pulmonary endothelium(7). We previously showed that BMP9 protects human pulmonary artery endothelial cells (hPAECs) against excessive permeability induced by TNF, LPS or thrombin(8). Moreover, administration of recombinant BMP9 protected mice against Evans Blue extravasation in the lung following intraperitoneal LPS challenge(8). Recently, adenoviral delivery of BMP9 has been shown to prevent retinal vascular permeability in diabetic mice(9). Despite such evidence suggesting that augmentation of BMP9 signaling might prevents endothelial hyperpermeability, the role of the endogenous BMP9 in the maintainence of endothelial barrier function, and the mechanisms involved, has not been investigated.

BMP9 is synthesized predominantly in the liver (10), circulates at levels that constitutively activate endothelial ALK1 signaling, and comprises the majority of plasma BMP activity (11). Heterozygous deleterious mutations in the *GDF2* gene (which encodes for BMP9) have been reported recently in patients with pulmonary arterial hypertension (PAH) (12-15), where they lead to reduced circulating levels of BMP9. Furthermore, reduced levels of plasma BMP9 are found in patients with cirrhosis and portopulmonary hypertension (16). Interestingy, increased vascular permeability is a well recognized feature of chronic cirrhosis.

Given the potential protective effect of BMP9 signaling in the pulmonary endothelium, we sought to determine whether endogenous BMP9 plays a role in protecting the pulmonary endothelium and whether inflammation regulates endogenous BMP9. We further explored the potential of exogenous BMP9 as a lung vascular protective agent in the setting of acute lung injury. Some of these results have been previously reported in the form of abstracts (17-19).

## MATERIALS AND METHODS

### Human Samples

Human plasma samples were obtained from a prospectively enrolled cohort of patients admitted to the adult medical intensive care unit (MICU) Registry of Critical Illness, and healthy human volunteers without known cardiopulmonary disease, in accordance with the Institutional Review Board–approved protocol at Brigham and Women’s Hospital as described previously (20-22). Written informed consent was obtained from all participants or their appropriate surrogates.

### Animal procedures

All procedures were carried out in accordance with the Home Office Animals (Scientific Procedures) Act 1986 and approved under Home Office Project Licences 80/2460 and 70/8850 to N.W.M.

### Murine endotoxemia studies

Mice were injected intraperitoneally with 2 mg/kg LPS or vehicle. After the length of time as specified in figure legends, mice were sacrificed using ketamine (100 mg/kg) and xylazine (10 mg/kg). Detailed tissue harvests and measurements can be found in an online data supplement.

### BMP9 ELISA

BMP9 ELISA was carried out as described previously (13, 16).

### Anti-BMP9 treatment in mice

Mice were injected intraperitoneally with 5 mg/kg anti-BMP9 antibody, or murine IgG2B isotype control on Day 0 and Day 2 and the lung vascular permeability was measured on Day 3. Separate groups of mice were injected intraperitoneally with 2 mg/kg LPS or PBS control on Day 2 and permeability was measured on Day 3 as a positive control. Half of the Lung tissues were harvested for measuring vascular permeability using Evans blue (EB) dye conjugated albumin as described previously (8), and the other half were inflated and fixed in formalin and processed into paraffin wax blocks for histological analysis.

### Expression and purification of pro-BMP9

Transfection and purification of pro-BMP9, consisting of the BMP9 prodomain non-covalently complexed with BMP9 growth factor domain (GF-domain), was achieved following previously described protocols (23, 24). Murine inhaled LPS model. Mice were injected intraperitoneally with either PBS or pro-BMP9 (at 1.5 μg/kg) 1 hour before being challenged with 20 μg/mouse LPS in PBS via the intranasal route. N=8 per group. After 24 hours, one lung was harvested for quantification of EB dye extravasation. The other lung was inflated and fixed in formalin and processed into paraffin wax blocks for histological analysis.

### Statistical analysis

All *in vitro* experiments were conducted at least three times, and representative images are shown. Data analysis was performed using GraphPad Prism 6. Results are shown as means ± SEM. Statistical significance was tested by two-tailed T-test or One-way ANOVA as indicated in the figure legends. Values of P < 0.05 were considered significant.

Further information can be found in an online data supplement, **Expanded Materials and Methods**. Microarray data have been deposited to Gene Expression Omnibus, with the accession number of GSE118353.

## RESULTS

### Inhibition of endogenous BMP9 increases lung vascular permeability and neutrophil extravasation

After neutralizing anti-BMP9 antibody was administered by intraperitoneal injection on day 0 and day 2, lung interstitial fluid accumulation was evaluated at day 3 using the EB dye extravasation assay (8)(Figure 1A). Mice treated with anti-BMP9 antibody exhibited significantly higher levels of interstitial fluid accumulation in their lungs compared with mice treated with isotype control IgG (Figure 1B). Remarkably, the magnitude of the anti-BMP9 antibody effect was similar to that observed in the lungs of mice exposed to intraperitoneal LPS (2 mg/kg, 24 hours) (Figure 1B). Histology of lung tissue demonstrated marked perivascular edema in mice exposed to either anti-BMP9 or LPS (Figure 1C, black arrows). Morphometry of arteries associated with terminal bronchioles confirmed acellular expansion of the adventitia, which correlated with the magnitude of EB dye accumulation in the lung (Figure 1D). Unexpectedly, anti-BMP9-treated mice showed a significant accumulation of neutrophils in the alveolar space, again similar to that observed in LPS-treated animals (Figure 1E). To confirm that anti-BMP9 antibody had indeed inhibited circulating BMP activity in mice, we tested plasma BMP9 activity using the readout of *ID1* gene induction in hPAECs as shown previously (10, 23). Indeed, plasma from anti-BMP9 treated animals showed reduced BMP activity, as evidenced by significantly lower *ID1* mRNA induction in hPAECs compared with plasma from IgG-treated controls (Figure 1F).

**Figure 1.**
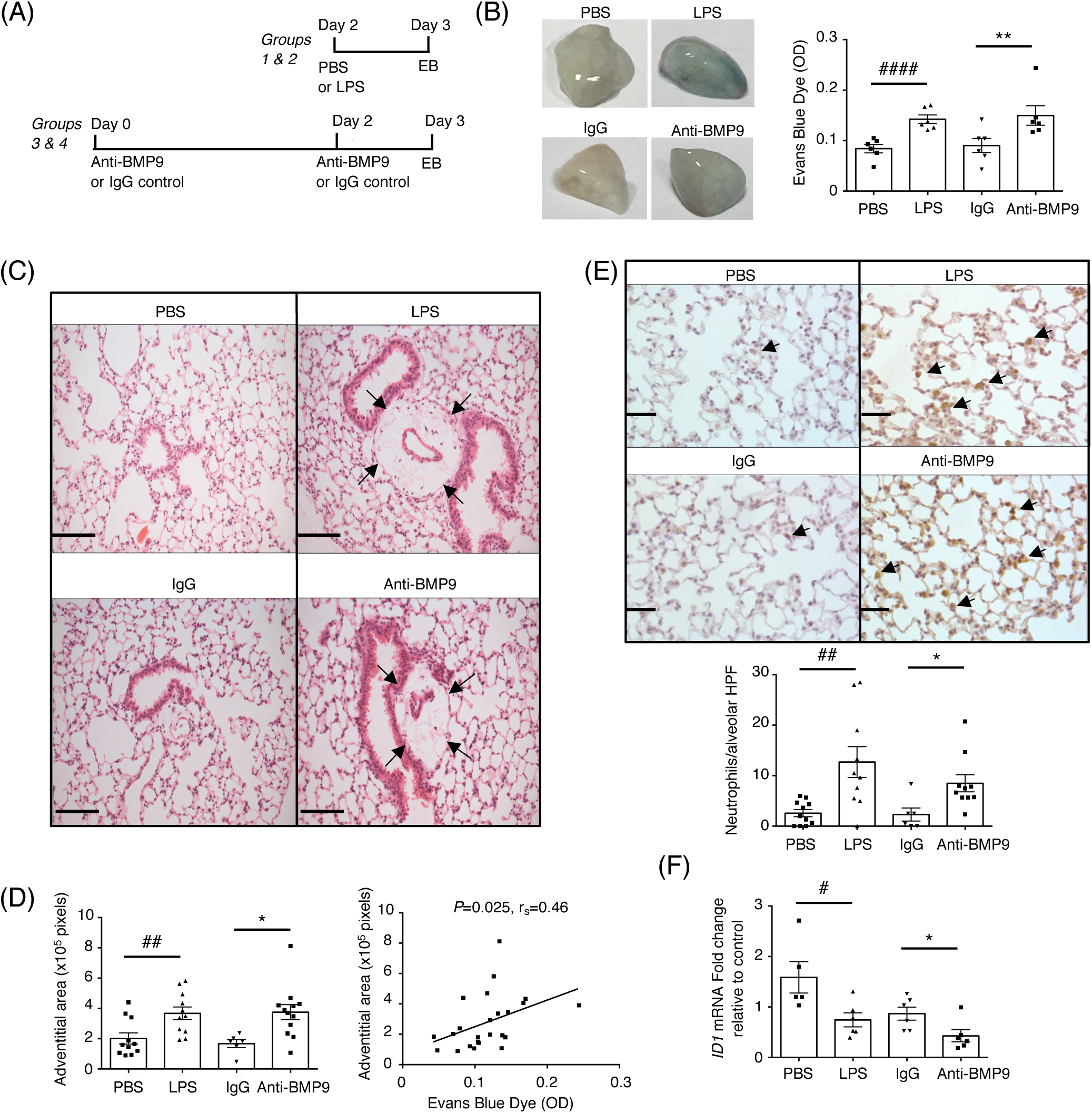
Neutralizing endogenous BMP9 results in lung vascular leak and neutrophil extravasation. (A) A schematic diagram indicating the treatment regimen. (B) Inhibiting endogenous BMP9 activity leads to lung vascular leak. Left, representative images of the lungs, showing the Evans Blue (EB)-stained lungs from LPS and anti-BMP9 treated animals. Right, quantification of EB content in the lungs. Data shown as means ± SEM, paired t-test with siblings from the same family as a pair, **, P ≤ 0.01; ####, P ≤ 0.0001. N=6 per group. (C&D) Anti-BMP9 treatment leads to an increase in the perivascular adventitial area, similar to LPS-treatment (black arrows). (C) Representative pictures of H&E stained lung section. Scale bars = 100 µM. (D) Adventitial area in 20 random high-power field (HPF), with its correlation to the EB content in the lung shown on the right; Spearman’s correlation test. (E) Anti-BMP9 treatment increases alveolar neutrophil counts, revealed by Myeloperoxidase staining. Scale bars = 50 µM. The counts were the mean of six random HPF per animal. For D&E, data are shown as means ± SEM, unpaired t-test, *, P < 0.05, ##, P ≤ 0.01. (F) BMP9 activity in plasma measured by *ID1*-gene induction in hPAECs. Serum-starved hPAECs were treated with 1% plasma samples for 1 hour before cells were harvested for RT-qPCR analysis of *ID1* gene induction. The operator was blinded to the treatment samples. Data are shown as means ± SEM, paired t-test, * and #, P < 0.05, N=6 per group.

### BMP9 signaling regulates key pathways involved in endothelial cell membrane integrity and permeability

To elucidate potential mechanisms by which BMP9 might act as an endothelial protective factor, we performed microarray analysis of differential gene expression in hPAECs to investigate global gene transcripts and pathways regulated by BMP9. We used the physiologically relevant form, prodomain bound BMP9 (pro-BMP9), at a concentration representative of levels measured in healthy human plasma, 0.4 ng/ml GF-domain (16, 25), and exposed hPAECs to pro-BMP9 for 5 hours. Using a threshold adjusted *P*-value of 0.05 as a cutoff, BMP9 upregulated the expression of 27 transcripts and downregulated 73 transcripts. However, the continuum changes in the adjusted P-values for both up- and down-regulated genes indicate that more transcripts are likely to be regulated than those passing this threshold (Figure 2A). Pathway analysis showed that BMP9-regulated pathways include TGFβ signaling, cytokine-cytokine receptor interaction and Rap1 signaling (see Figure E1 and Table E1&2 in the online data supplement). Gene ontology analysis revealed that BMP9 regulated genes are highly enriched in the plasma membrane and extracellular space (Figure 2B). As expected, BMP9 increased the expression of *BMPR2* and *ID1* (3) (Figure 2A). Of the genes known to regulate endothelial permeability, pro-BMP9 treatment down-regulated *AQP1* (encoding aquaporin-1) and *KDR* (encoding VEGFR2), and up-regulated *TEK* (encoding Tie2) (Figure 2A). Up-regulation of *TEK* and down-regulation of *AQP1* and *KDR* by BMP9-signaling in hPAECs were confirmed by RT-qPCR (Figure 2C).

**Figure 2.**
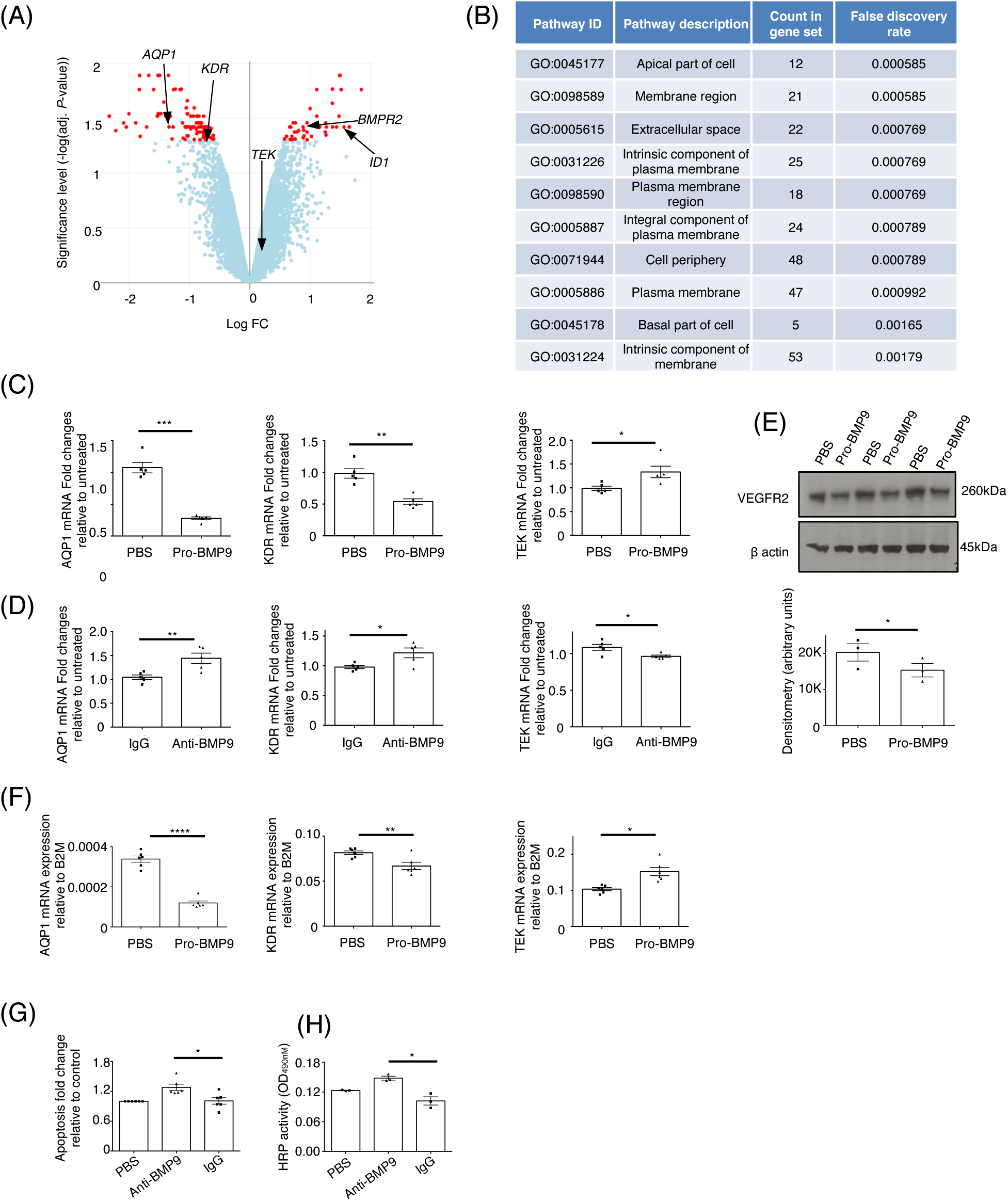
BMP9 signaling regulates genes involved in endothelial cell integrity. (A) A volcano plot of microarray transcriptional analysis of BMP9 regulated genes. Serum-starved hPAECs were treated with 0.4 ng/ml (GF-domain concentration) pro-BMP9 for 5 hours before cells were harvested for microarray analysis. Four independent hPAEC lines were used. Adjusted P-values of less than 0.05 are shown in red. (B) Microarray results were subject to Cellular Component GO analysis pathway using STRING Sever (https://string-db.org/). (C) Validation of microarray results using RT-qPCR. BMP9 signaling regulates mRNA expression of *AQP1* (Aquaporin1), *KDR* (VEGFR2) and *TEK* (Tie2) in hPAECs. N=5. (D) Change of gene expression in hPAECs after inhibition of BMP9 activity in FBS with a neutralizing anti-BMP9 antibody. PAECs were grown in endothelial basal medium with 2% FBS, treated with IgG control or anti-BMP9 antibody (both at 20 μg/ml) for 3 hours (for *TEK*) or 5 hours (for *AQP1* and *KDR*) before cells were harvested for RNA extraction and qPCR analysis. N=5. (E) BMP9-treatment suppresses VEGFR2 total proteins. Serum-starved hPAECs were treated with pro-BMP9 (0.4 ng/ml GF-domain concentration) for 5 hours. Three independent treatments were run on the same western blot and quantified using ImageJ. (F) BMP9 regulates *AQP1, KDR* and *TEK* expressions in human pulmonary microvascular endothelial cells (hPMECs). (G) In hPMECs, anti-BMP9 treatment leads to enhanced apoptosis measured using Caspase 3/7 Glo assay. (H) Anti-BMP9 treatment in hPMECs causes enhanced monolayer permeability measured by HRP-transwell assay as described previously(8). For all the panels, means ± SEM are shown, two-tailed, paired t-test, *, P < 0.05; **, P ≤ 0.01; ***, P ≤ 0.001. ****, P ≤ 0.0001.

In addition to adding exogenous BMP9, we performed the converse study: neutralization of endogenous BMP9 in serum. Selective removal of BMP9 from FBS-containing endothelial growth media using neutralizing anti-BMP9 antibody generated reciprocal changes in gene expression compared to IgG-treatment controls: a significant increase in *AQP1* and *KDR* expression and a reduction in *TEK* expression (Figure 2D), confirming that endogenous serum BMP9 regulates these genes even in the presence of other serum factors. To confirm the regulation of VEGFR2 at the protein level, immunoblotting was perfomed. BMP9 consistently suppressed VEGFR2 protein expression at both 5 hours and 8 hours post pro-BMP9 treatment (Figure 2E, only 5 hour data shown). AQP1 expression in cultured PAECs was too low to be accurately detected by western blotting.

Given that the lung microvascular endothelium rather than the conduit artery endothelium is involved in lung hyperpermeability, we further examined responses in human pulmonary microvascular endothelial cells (hPMECs). Similar to hPAECs, BMP9 induced robust changes in gene expression in hPMECs, exemplified by the induction of *ID1* and *BMPR2* expression (Figure E2). We also confirmed that BMP9 treatment suppresses *AQP1* and *KDR* expression, and induces *TEK* expression in hPMECs (Figure 2F). Functionally, anti-BMP9 treatment in PMECs led to enhanced apoptosis (Figure 2G) and increased permeability (Figure 2H) compared with IgG-treated controls.

Taken together, these data strongly support a role of BMP9 signaling in regulating endothelial cell membrane integrity and barrier function.

### Exogenous BMP9 protects mice from acute lung injury in response to inhaled LPS

Acute inhaled administration of LPS initiates epithelial cell damage and causes lung vascular hyperpermeability and injury in mice. To further explore the potential therapeutic value of BMP9 (8), we questioned whether administration of exogenous BMP9 prevents lung injury in this murine model. As expected, inhalation of LPS led to pulmonary inflammation and congestion (Figure 3A) and increased lung vascular permeability measured by extravasation of EB dye (Figure 3B). All of these features were completely prevented by pre-treatment of mice with BMP9. The degree of lung injury was scored following the recommendation of the American Thoracic Society workshop report (26) (Figure 3C), and the number of neutrophils extravasated into alveoli were counted (Figure 3D). Supplementation of BMP9 resulted in a complete protection against LPS-induced acute lung injury and neutrophil extravasation. Interestingly, in this model, plasma levels of neutrophil elastase were also elevated following LPS treatment and this increase was prevented in the BMP9 pre-treated animals (Figure 3E), suggesting an anti-inflammatory role of BMP9. To confirm target engagement, we investigated the changes in BMP9-regulated genes in the lung. As expected, administration of BMP9 led to an enhancement in BMP signaling, as evidenced by the elevated mRNA expression of the BMP9 target genes *ID1* and *BMPR2* (Figure 3F&G). Of note, consistent with the results of *in vitro* studies (Figure 2), the expression of *TEK* (Tie2) was significantly reduced following LPS challenge, and restored by BMP9 treatment (Figure 3H). Furthermore, LPS treatment led to a 2-fold increase in expression of *KDR* (VEGFR2), which was completely prevented by the pre-treatment with BMP9 (Figure 3I).

**Figure 3.**
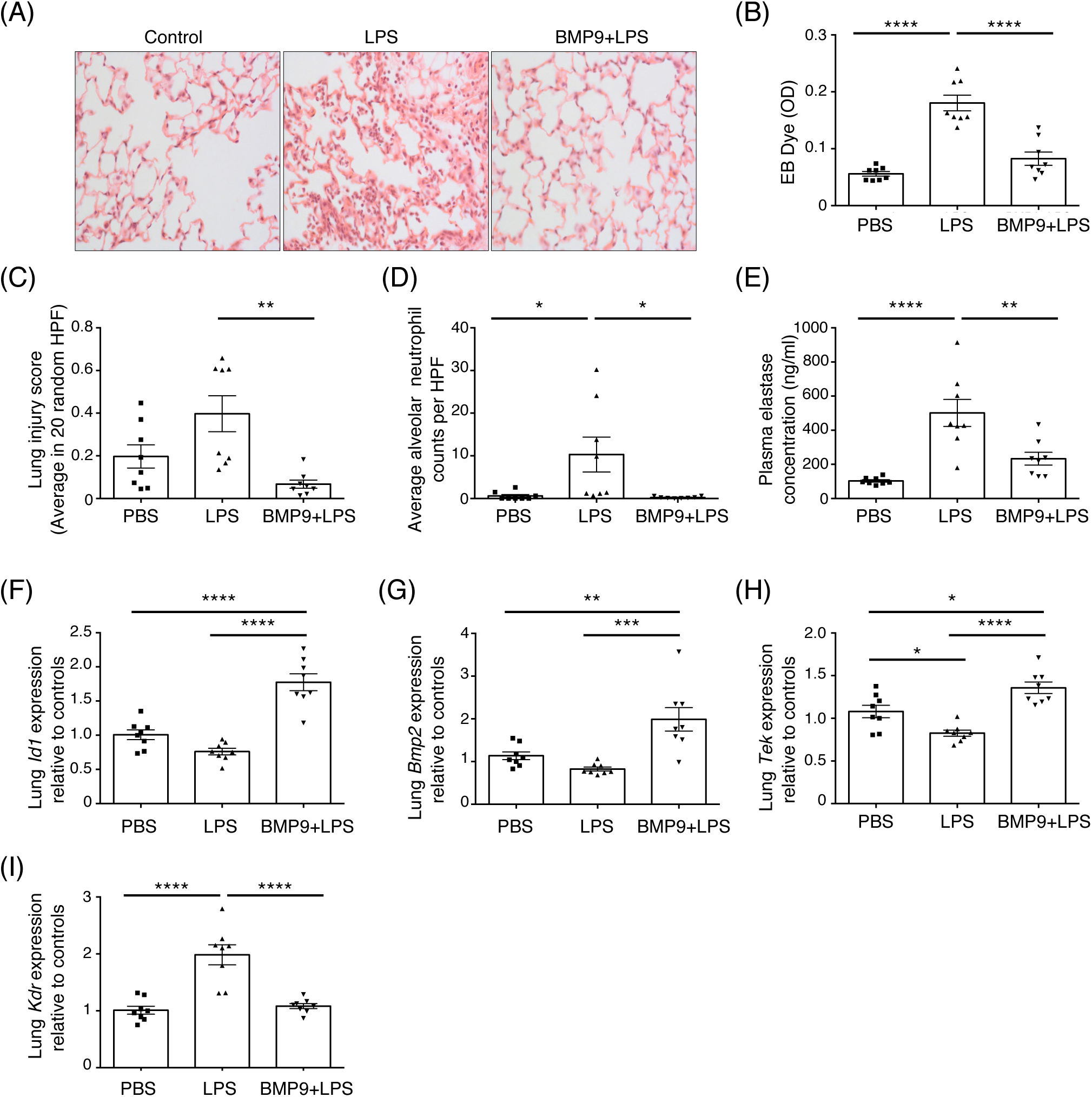
BMP9 prevents vascular leak and lung injury in inhaled LPS-challenged mice involves *TEK* and *KDR*. (A) Representative images of H&E-stained lung tissues. Mice were challenged intranasally with LPS (at 20 μg/mouse) for 24 hours before lungs were harvested for immunohistological examination. (B) BMP9 prevented vascular leak measured by EB retained in the lungs. (C) BMP9 protected acute lung injury induced by inhaled LPS. Lung injury scores were based on 20 HPF per animal as per protocol from the American Thoracic Society Workshop report (26). (D) Administration of BMP9 prevented the extravasation of neutrophils into the alveolar space. Neutrophils were counted from the H&E-stained slides based on the shape of the cells and nuclei. (E) Administration of BMP9 prevented the increase of plasma elastase after inhaled LPS-challenge. (F-I) Lung mRNA expression measured by qPCR. RPL32 was used as the house-keeping gene. The operator was blinded to the treatment in this experiment. For all panels, means ± SEM are shown, one-way ANOVA, Tukey’s post test. *, P < 0.05; **, P ≤ 0.01; ***, P ≤ 0.001; ****, P ≤ 0.0001.

### Plasma BMP9 is suppressed during endotoxemia

Given that both depletion or supplementation of endogenous BMP9 impacts lung vascular permeability, we questioned whether circulating BMP9 levels are reduced per se in the setting of systemic inflammation in humans and mice. Eighteen hours after intraperitoneal LPS administration, mice exhibited a systemic inflammatory response, as evidenced by a reduction in platelet numbers and an increase in alveolar neutrophil counts (Figure E3 A&B). Concentrations of neutrophil elastase were also significantly elevated in both the plasma and bronchoalveolar lavage fluid (Figure E3 C&D). As hypothesized, circulating levels of BMP9 were markedly decreased in this murine model (Figure 4A).

**Figure 4.**
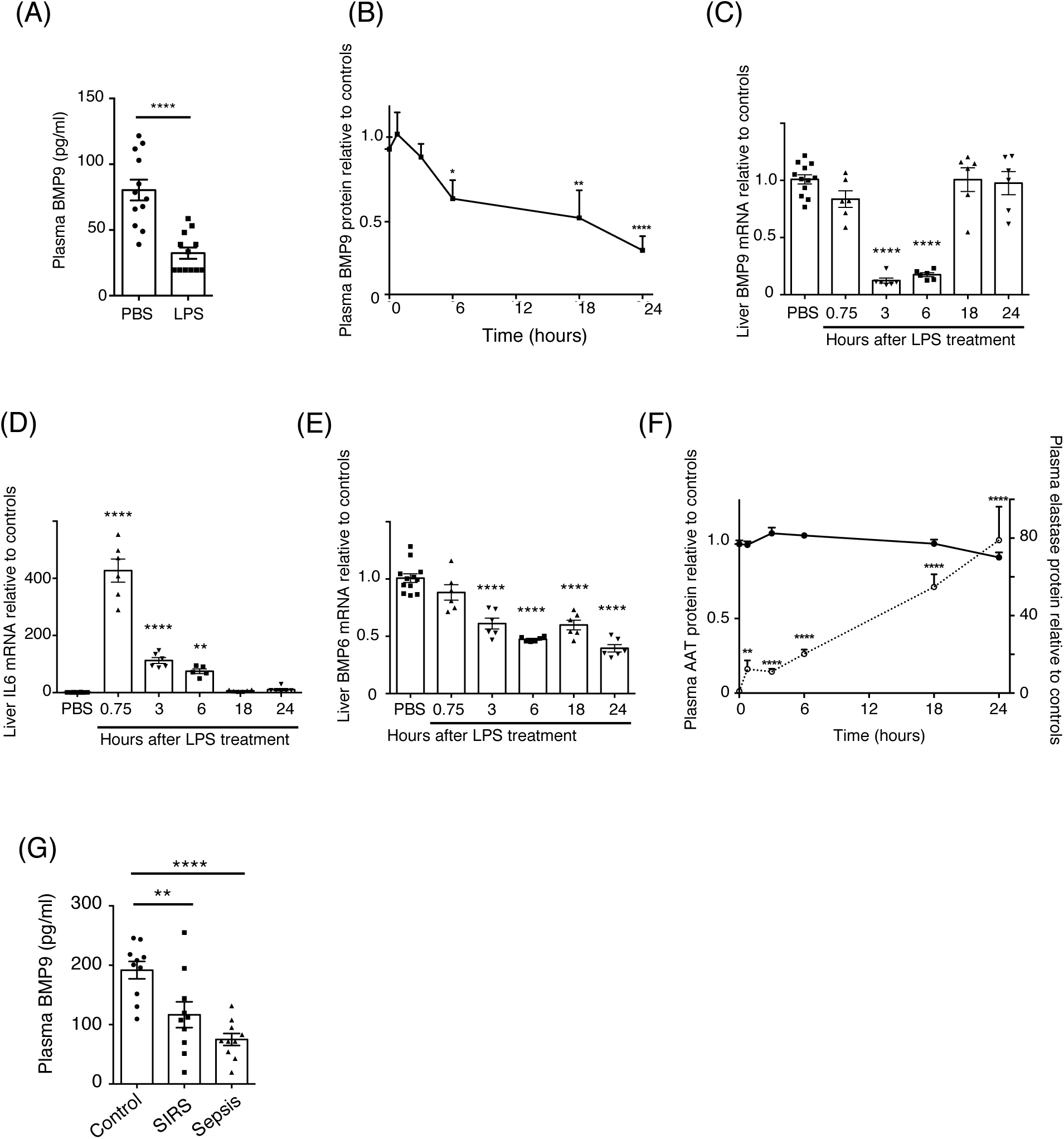
Endogenous BMP9 is reduced in endotoxemia mice and sepsis patients. *(A) Circulating BMP9 levels are significantly reduced in murine endotoxemia model*. Mice were treated with 2 mg/kg LPS intraperitoneally for 18 hours before plasma were taken for BMP9 measurement. Data are shown as means ± SEM. Two-tailed, unpaired t-test, ****, P ≤ 0.0001. *(B) Dynamic changes in circulating BMP9 after LPS-induced inflammation*. Mice were treated with 2 mg/kg LPS intraperitoneally and sacrificed at 0, 0.75, 3, 6, 18 and 24 hours. N=6 per group. Three animals were treated with PBS at each time point and used as controls. Concentrations of BMP9 in plasma were measured by ELISA, normalized to controls. *(C-E) Dynamic changes of liver mRNA expression relative to controls after LPS-challenge*. Data were analyzed using the ΔΔCt method, using RPL32 as the house keeping gene. *(F) Changes in plasma elastase (open circle, dashed line) and AAT (filled circle, solid line) protein levels relative to controls during endotoxemia*. The actual control value for AAT is 3.44 ± 0.09 mg/ml, for elastase is 128.6 ± 26.8 ng/ml. In all measurements, means ± SEM are shown, one-way ANOVA, Dunnett’s post test against PBS controls, *, *P* < 0.05; **, *P* ≤ 0.01; ****, *P* ≤ 0.0001. *(G) Plasma BMP9 concentrations from SIRS and sepsis patients are significantly lower than those from healthy controls*. One-way ANOVA, Dunnett’s post test. **, *P* ≤ 0.01, ****, *P* ≤ 0.0001.

To further delineate the changes in the endogenous BMP9 during inflammation, we performed a time-course study to track the mRNA and protein changes in BMP9 following the onset of endotoxemia in mice. Significant decreases in plasma BMP9 levels were detected from 6 hours after LPS exposure and BMP9 levels continued to fall at the 24-hour time point (Figure 4B). Hepatic Bmp9 mRNA levels were suppressed to ∼20% of control values at 3 hours post LPS exposure (Figure 4C), but returned to control values by 18 hours, despite the continued reduction in plasma BMP9 levels at these time points. As comparators, we also measured liver mRNA levels for IL-6 to monitor the inflammatory response, and BMP6, which is another BMP known to be expressed in the liver (27). There was a sharp increase in Il6 mRNA at 45 minutes after LPS administration, which fell at 3 hours and returned to baseline levels at 18 hours (Figure 4D). Bmp6 mRNA was reduced to about 60% of the control levels by 3 hours and remained suppressed throughout the 24 hour period (Figure 4E). These comparators confirm that the rapid loss and the subsequent recovery of Bmp9 mRNA levels are unique for BMP9 and not due to the global suppression and recovery of mRNA synthesis in the liver.

As the continued reduction in plasma BMP9 during mouse endotoxemia could not be explained fully by the changes in hepatic *Bmp9* mRNA expression alone, we investigated whether BMP9 might also be degraded by plasma proteases. Inflammation leads to the activation of neutrophils, which release large amounts of proteases, especially elastase (28); we therefore investigated whether LPS challenge causes changes in circulating neutrophil elastase (NE) levels. Compared with PBS-treated control animals, administration of LPS caused a 10-fold increase in plasma elastase concentration at 45 minutes, and an 80-fold increase at 24 hours (Figure 4F). Because α1-antitrypsin (AAT) is the major NE inhibitor in plasma (29), we also measured AAT liver mRNA and plasma protein levels. AAT mRNA levels were largely unchanged over the first 6 hours but decreased to about 50% of controls at 18 and 24 hours (Figure E4 A). Using an ELISA that specifically detects the native and active form of AAT (Figure E4 B) (30), we observed that plasma levels of active AAT were largely unchanged throughout the 24-hour time course following LPS challenge (Figure 4F). This indicated that the 80-fold increase of NE levels were not counteracted by a similar fold increase in this endogenous inhibitor, leading to an imbalance favoring heightened elastase activity during endotoxemia. To examine whether circulating BMP9 levels are decreased in patients with systemic inflammatory response syndrome (SIRS) and sepsis, plasma BMP9 levels were measured in 10 patients with SIRS, 10 patients with sepsis, all sampled within 72 hours of admission to the intensive care unit, and 10 age- and sex-matched healthy controls. The clinical characteristics and the demographics of the subjects are summarized in Table E3 and are notable for the presence of positive microbiology cultures and an increased reliance upon vasopressors for blood pressure support in the sepsis patients compared with the other groups. Importantly, plasma BMP9 levels were significantly reduced in patients with SIRS and further reduced in sepsis, compared with healthy controls (Figure 4G).

### BMP9 is a substrate for neutrophil elastase

Finally we sought to confirm whether BMP9 can be cleaved by NE. Using purified recombinant proteins, the circulating form of prodomain-bound BMP9 can be cleaved efficiently by NE, despite being highly resistant to trypsin digestion (Figure 5A). Next, we questioned whether NE in the plasma could contribute to BMP9 cleavage, since primed circulating neutrophils (with increased capacity for systemic degranulation) were identified in ARDS patients(31), activated neutrophils release a number of proteases upon degranulation(28) and significantly higher levels of NE in plasma from endotoximic mice (Figure 4F). Purified human peripheral blood neutrophils were activated *in vitro* to degranulate and release proteases into the culture supernatant as described previously (28) (Figure 5B). Pro-BMP9 was incubated with supernatants derived from activated neutrophils, in the presence or absence of a panel of protease inhibitors (PI). As shown in Figure 5 C&D, activated neutrophil supernatants were effective at cleaving BMP9 and this process was blocked by AAT and sivelestat, a selective NE inhibitor. In contrast, the chelating agent EDTA did not confer any inhibition, suggesting that metalloproteases do not play a role in BMP9 cleavage. Taken together these findings suggest that NE contributes to the cleavage of BMP9 in the settting of inflammation.

**Figure 5.**
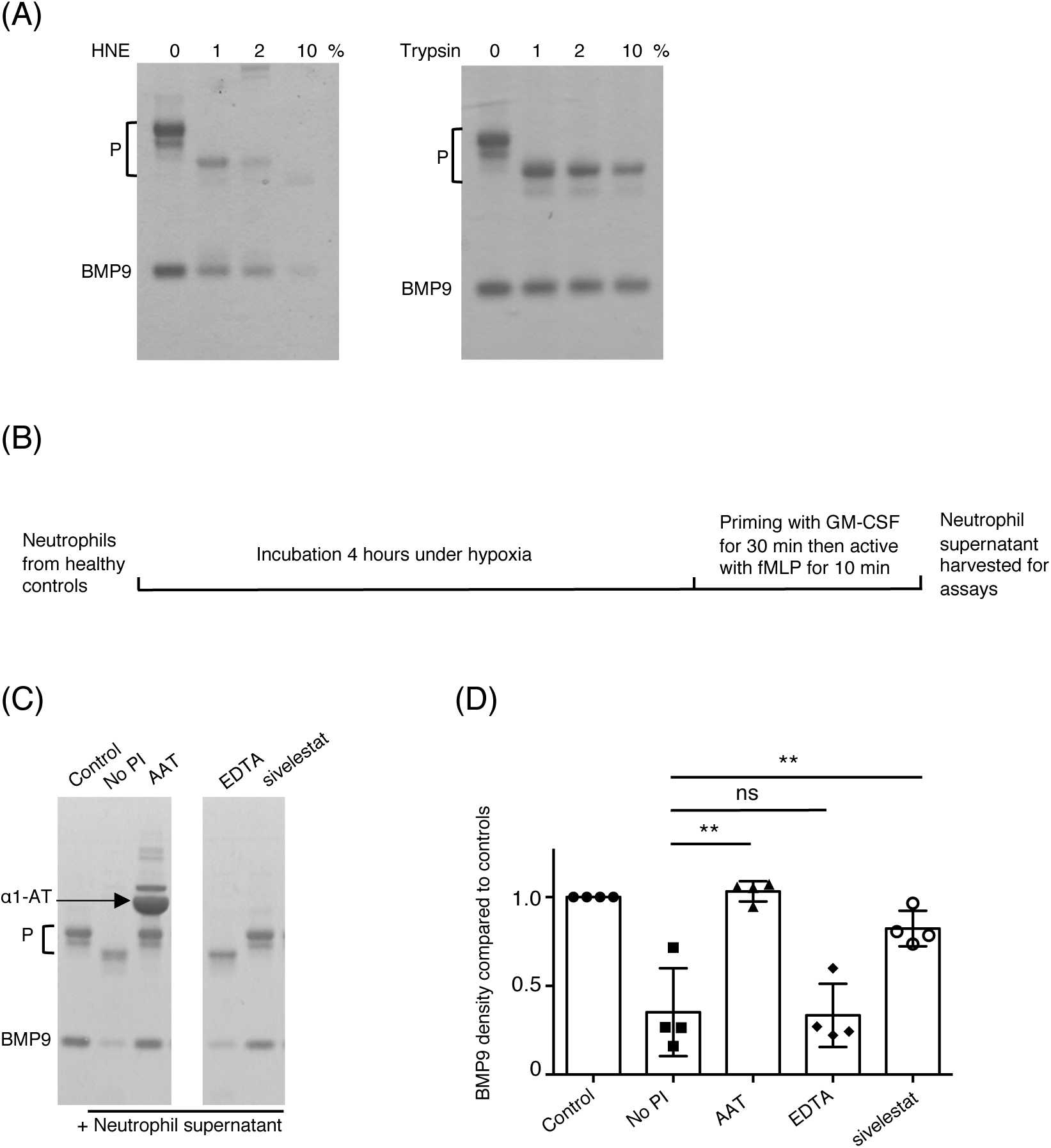
BMP9 is a substrate for neutrophil elastase. *(A) BMP9 is a direct substrate of elastase*. Purified pro-BMP9 was incubated with recombinant human neutrophil elastase (HNE) or trypsin at indicated concentration (% w/w) in PBS overnight and the mixture was fractionated by SDS-PAGE under reducing conditions. P: prodomain. *(B) A schematic diagram illustrating the generation of supernatants from activated neutrophils. Neutrophils* were isolated from the peripheral blood of the healthy volunteers, incubated under hypoxia for 4 hours before priming with GM-CSF and activation with fMLP as described previously (28). *(C&D) Neutrophil elastase is the major protease cleaving pro-BMP9 in the activated neutrophil supernatant*. Pro-BMP9 was incubated with the supernatant from activated neutrophils in the presence or absence of a panel of protease inhibitors overnight and the mixture fractionated by SDS-PAGE under reducing conditions. BMP9-cleavage activity can be inhibited only by serine protease inhibitor AAT, and elastase-specific inhibitor sivelestat, not by EDTA which inhibits metalloproteases. A representative gel from 4 independent experiments is shown in (C, two parts of the same gels are shown) and the quantification of BMP9 bands from 4 experiments in (D). Means ± SEM are shown, paired t-test. *, *P* < 0.05; **, *P* ≤ 0.01; ns, not significant.

## DISCUSSION

In this study we provide evidence that endogenous BMP9 is an important protective factor in the pulmonary vascular endothelium, which is significantly down-regulated during inflammation. Selective inhibition of circulating BMP9 using a neutralizing antibody alone was sufficient to induce heightened lung vascular leak. Such a finding is consistent with reports that ALK1-Fc, a ligand trap of BMP9, causes peripheral edema as a common side effect in clinical trials (32, 33). In addition, the major endothelial type 2 receptor for BMP9, BMPR-II, has been shown to play a role in endothelial integrity. Loss of BMPR-II promotes endothelial permeability and contributes to the development of PAH (8, 34). It is tempting to speculate that the pulmonary circulation is particularly dependent of BMP9/ALK1 signaling to maintain barrier function since the ALK1 receptor is particularly highly expressed on lung vascular endothelium (7).

We show here for the first time that the circulating form of BMP9, at physiologically relevant concentrations, regulates the transcription of gene sets highly associated with the plasma membrane and the extracellular space, including three receptors controlling critical pathways involved in endothelial permeability. Importantly, we show that *in vivo*, the protection by BMP9 against lung vascular leak in the murine inhaled LPS model was associated with the preservation of *TEK* and *KDR* expression in the LPS-exposed lung. The effect of VEGF signaling via VEGFR2 to induce vascular leak has been extensively studied (35). Tie2 mediates the signaling of angiopoietin 1 (Ang1), and Ang1 prevents vascular leak via multiple mechanisms, including inhibition of VEGF signaling (36). Consistent with a role for Tie2, a recent study identified increased expression of angiopoietin 2 (Ang2) in the retina of neonatal mice receiving anti-BMP9 antibodies (37). *AQP1* regulates osmotically driven water transport across microvessels in adult lungs and facilitates hydrostatically driven lung edema (38). Decreased pulmonary vascular permeability has been described in *AQP1*-null humans (39) and *AQP1* expression is increased in the capillary endothelium of alveoli from patients with ARDS (40).

It is interesting to note the inverse correlation of circulating BMP9 and elastase concentrations in the onset of endotoxemia in mice, and that BMP9 is a direct substrate of NE *in vitro* despite it being highly resistant to trypsin digestion. Further experiments are needed to show BMP9 cleavage by NE *in vivo*. This could be challenging because we showed here that NE is a major but not the only serine protease released by neutrophils that has the ability to cleave BMP9 (Figure 5C&D), therefore elastase inhibition alone may not be enough to rescue BMP9 levels in circulation. A direct detection of elastase-cleaved BMP9 fragments *in vivo* would be more informative, however, this is difficult due to the presence of very low concentrations of BMP9 in the circulation (200-400 pg/ml).

We previously reported that BMP9 enhances LPS-induced leukocyte recruitment to the vascular endothelium(41). However, this effect was observed with higher concentrations of BMP9, that are likely activating the ALK2 receptor(42). The present study used lower concentrations of BMP9, and the data in this study are consistent with the results reported by Burton et al (34) and Long et al(8). Since BMP9 can signal through both the high affinity receptor ALK1, as well as low affinity receptor ALK2, our overall results suggest that restoration of BMP9 levels, around the physiological concentration range, will cause BMP9 to signal through the ALK1-mediated pathway and exert beneficial anti-inflammatory and endothelial-protective effects.

Microvascular leak has now been recognized as a major contributor to septic shock and is associated with increased morbidity and mortality; as yet, there is no pharmacological drug currently available that targets this process (43). Restoration of endothelial integrity has been shown to increase survival in three different animal models of systemic inflammation (44). Our study identifies a new and unexpected essential role for endogenous BMP9 in the maintenance of endothelial barrier function under physiological conditions, and demonstrates that circulating BMP9 levels are reduced in sepsis patients and in a murine endotoxemia model. Importantly, administration of BMP9 protects against lung vascular leak in the murine acute lung injury model. Taken together, these findings support the exploration of BMP9 as a biomarker as well as a potential therapy for the prevention of vascular permeability and lung injury associated with sepsis and ARDS.

## ACKNOWLEDGEMENTS

We are grateful to Dr James Thaventhiran for providing training on the inhaled LPS procedure, MICU Registry members Laura Fredenburgh, Paul Dieffenbach, Samuel Ash at Brigham and Women’s Hospital for collecting human plasma samples. We also thank the support from Cambridge Genomic Service, in particular Dr. Emily Clemente and Dr. Julien Bauer, for the microarray data analysis.

## Notes

Conflict of Interest: WL and PDU are founders and consultants to Morphogen-IX. NWM is a founder and CEO of Morphogen-IX.

Funding: This work was supported by the following grants: British Heart Foundation grants (PG/17/1/32532 to WL and NWM, PG/17/58/33134 to WL, NWM and CS, CH/09/001/25945 and RG/13/4/30107 to NWM), MRC grant (MR/K020919/1 to NWM), British Lung Foundatin grant (BLF COPD10/5 to AMC), and National Institutes of Health Grants (P01-HL108801 to RMB, R01-HL131910 to PBY and R42-HL132742 to PBY). KML was supported by a Wellcome Trust Clinical Research Fellowship (102706/Z/13/Z). Infrastructre support was provided by the Cambridge NIHR Biomedical Research Centre.

### Competing Interest Statement

Prof Morrell, and Drs Wei Li and Paul Upton are co-founders and scientific advisers to Morphogen-IX.

