## Supplementary material for "Circulating BMP9 protects the pulmonary endothelium during inflammation-induced lung injury in mice"

**Endogenous BMP9 protects pulmonary vascular endothelium and is down-regulated during inflammation**

### ONLINE DATA SUPPLEMENT:

**Expanded materials and methods**

**Table E1.** Summary of pathway analysis of BMP9 regulated protein-protein interaction network

**Table E2.** Summary of enriched pathways in hPAECs from pathway analysis of BMP9 regulated protein-protein interaction network

**Table E3.** Patient demographics and clinical parameters

**Figure E1.** BMP9 regulated genes are highly enriched on plasma membrane or extracellular space.

**Figure E2.** BMP9 signaling in hPMECs.

**Figure E3.** Murine model of endotoxemia induced by acute intraperitoneal LPS challenge.

**Figure E4.** The regulation of endogenous BMP9 during endotoxemia

**References**

### Expanded materials and methods

#### *Materials:*

Male C57BL/6 mice (12 weeks old, ~25-30g) were purchased from Charles River and used for all *in vivo* studies. Mouse elastase ELISA kit was purchased from Cloud-Clone Corp (Cat No. SEA181Mu). Mouse α1-antitrypsin ELISA kit was purchased from GenWay Biotech Inc (Cat No. GWB-DB430D). Both mouse monoclonal (Cat No. MAB3209) and biotinylated goat polyclonal (Cat No. BAF3209) anti-BMP9 antibodies, and IgG2B (MAB004) were purchased from R&D Systems. Endothelial cell basal media (EBM-2) were purchased from Lonza or PromoCell; and supplemented with a Bullet kit (Lonza) to obtain endothelial growth medium (EGM-2). All plasmid and RNA purification kits were purchased from Qiagen. LPS was from Sigma Aldrich (Cat No. L2630). Total blood count was carried out using the scil Vet ABC ™Hematology Analyzer (Woodley Equipment Company LTD). Recombinant human neutrophil elastase was purchased from AppliChem.

***Neutrophil elastase cleavage assay.*** Neutrophils were isolated from peripheral blood and activated *in vitro* under hypoxia as described previously(1). Five μg of recombinant pro-BMP9 was incubated at room temperature overnight, in a final volume of 20 μl in PBS, containing 5 μl of activated neutrophil supernatant (generated as described previously (1)) without or with protease inhibitors. The mixture was separated on a 12% SDS-PAGE (NuPAGE, Invitrogen) under reducing conditions and visualized by Coomassie Blue staining. The band intensities of BMP9 growth factor domain were quantified using Image J.

***Murine endotoxemia studies.*** Mice were injected intraperitoneally with 2 mg/kg LPS or vehicle. After the length of time as specified in figure legends, mice were sacrificed using ketamine and xylazine. Bronchoalveolar lavage fluid (BALF) was collected by injecting and recovering 1 ml PBS via trachea. Blood for plasma preparation was taken via the inferior vena cava. Liver and right lung were snap frozen for RNA extraction while the left lung was inflated with a 1:1 mixture of saline and O.C.T. compound (Sakura, Zoeterwoude, Netherlands), fixed with 4% paraformaldehyde in PBS before dehydration and paraffin embedding for immunohistochemistry.

***Measuring gene expression in mouse liver*****.** Liver total mRNA was extracted using Qiagen miRNeasy Mini kit following manufacturer's instruction. After measuring the mRNA concentration on a Nanodrop Spectrometer, 1 ng was included in the reverse transcription reaction using High-Capacity cDNA Reverse Transcription Kit (Applied Biosystems) following the manufacturer's instruction. Quantitative PCR (qPCR) was carried out in 384 well plate on a Quantstudio 6™ Real-Time thermal cycler (Applied Biosystems) and the relative expression of target mRNA was normalized to Ribosomal Protein L32 (*Rpl32*) using the ΔΔCT method(2), expressed as the fold-changes relative to PBS treated controls. The following forward and reverse primers were used to amplify the corresponding mouse genes: α1-AT: 5' GGCCATACCCATGTCTATCC-3'; 5' TTCACCACTTTTCCCATGAA-3', *Bmp9*: 5'-ACAACGGACAAATCGTCTACG-3'; 5'- AGGATGTGCTTCTGAAAGGGG-3'. QuantiTect primers (Qiagen) were used for mouse *Bmp6* and *Il6*.

***Haematoxylin and eosin (H&E) staining and lung injury scoring.*** Four μm mouse lung sections were H&E stained and examined by light microscopy. Lung injury was scored following the published guidelines (3). Mice were immunostained using rabbit-polyclonal anti-myeloperoxidase (DakoCytomation, UK), labelled using immunoperoxidase (Vectastain Elite, Vector Laboratories) and 3,3′-DAB to create a brown coloured reaction product as previously described (4). All immunohistology scorings were carried out in a blinded manner.

***PAEC signaling assay for evaluating plasma BMP9 activity.*** Human PAECs were grown to ~80% confluence, serum-starved overnight in EGM-2 containing 0.1% FBS before treating with 1% mouse plasma. After 1 hour, cells were snap-frozen to stop the signaling and harvested for mRNA analysis. RNA extraction, reverse transcription and qPCR analysis were carried out as described above. The following primers were used for the qPCR reactions: human *ID1*: 5'-CTGCTCTACGACATGAACGGC-3', 5'-TGACGTGCTGGAGAATCTCCA-3'; human β2 microglobulin (*B2M*): 5'-CTCGCGCTACTCTCTCTTTCT-3', 5'-CATTCTCTGCTGGATGACGTG-3'.

***Pro-BMP9 signaling in hPAECs and microarray.*** For the microarray experiment, 4 different lines of hPAECs were serum-starved overnight in EGM-2 containing 0.1% FBS before being treated with 0.4 ng/ml pro-BMP9 for 5 hours. Cells were snap-frozen and harvested for mRNA extraction and microarray analysis. Microarray experiments were performed at Cambridge Genomic Services, using a human Gene 2.1 ST Array Plate (Affymetrix, Wooburn Green, UK) in combination with WT PLUS amplification kit (Affymetrix) according to the manufacturer's instructions. Following data processing using package Oligo in R (5), normalization was carried out using Robust Multichip Analysis (RMA)(6), and comparisons were performed using the limma package (7). The results were corrected for multiple testing using False Discovery Rate (FDR). Protein-protein interaction (PPI) and enriched pathway analysis were performed using STRING (8).

Validation of the target genes identified by microarray were carried out by repeating the hPAEC treatment as above and extracted RNA were subject to RT-qPCR analysis. The following primers were used for the qPCR reactions: human *AQP1*: 5'TCTCAGGCATCACCTCCTCC-3', 5'CGAGTTCACACCATCAGCCA-3'; human *KDR*: 5'GATGCAGGAAACTACACGGTCA-3', 5'TCCATAGGCGAGATCAAGGCT-3'; human *TEK*: 5'GAAACATCCCTCACCTGCATTG-3', 5'TTTCGCCCATTCTCTGGTCA-3'. Microarray data have been deposited to Gene Expression Omnibus, with the accession number of GSE118353.

***Anti-BMP9 treatment of hPAECs for qPCR.*** Human PAECs were cultured in EBM-2 containing 2% FBS for at least 20 h before treatment with 20 μg/ml anti-BMP9 antibody (MAB3209, R&D systems) or 20 μg/ml IgG2B (MAB004, R&D systems) for 3 or 5 hours. Cells were snap-frozen to stop the signaling and harvested for mRNA analysis. RNA extraction, reverse transcription and qPCR analysis were carried out as described above.

**Apoptosis assay.** Apoptosis assay was performed using Caspase-Glo 3/7 Assay System (Promega, Cat. No. G8090). PMECs were seeded in triplicates at 10,000 cells per well in 96-well cell culture plates overnight in EGM-2 containing 10% FBS. On the following day, cells were incubated with 100 μl fresh endothelial basal medium (EBM-2) containing 2% FBS with different treatment reagents: antiBMP9 antibody (at 100 μg/ml), IgG (at 100 μg/ml), or vehicle control (0.5%BSA in PBS). After 5 hours, cells were harvested by adding 100 μl substrate reagent to each well. The plates were protected from light by foil and mixed at 400 rpm for 20 min. 100 μl supernatant was transferred to a 96-well white wall plate and luminescence changes were recorded in a microplate luminometer.

**Permeability assay.** PMECs were seeded at 1 x $10^5$ cells per trans-well insert (Costar) in EGM-2 with 10% FBS and incubated for 24 hours. On the next day, inserts were moved to a new chamber with 2 % FBS in EBM-2, and the upper chamber medium were changed to 2 % FBS in EBM-2 containing 20 nM HRP that was supplemented with anti-BMP9 antibody (at 100 μg/ml) or IgG (at 100 μg/ml) or an equal volume of medium. At 10 min, 15 μl medium from the lower chamber was collected and transferred to a 96 well plate. 150 μl buffer containing HRP substrate (o-phenylenediamine dihydrochloride) was added and the plate was read at 490 nm immediately.

**Table E1. Summary of pathway analysis of BMP9 regulated protein-protein interaction (PPI) network.** In the microarray analysis, we used N=4 hPAEC lines (isolated from four different individuals), and at a single time point, 5 hours. The continuum changes in the adjusted *P*-values (adj *P*) for differentially regulated genes indicate that the up and down regulated genes are likely to be more than those passing the adj *P* < 0.05 threshold, and the reasons for not reaching the adj *P* of 0.05 cut-off could be due to either the sample number was too small, or the peak time for the change is not at 5 hours. Therefore, we have performed the PPI network analysis using the gene set with adj *P*-value cut-off at 0.05, 0.06 and 0.08. The analysis was performed using STRING(8). The network stats summaries of all analyses are shown below, enriched pathways shown in Supplemental Table E2 and Network view of differentially regulated genes shown in Supplemental Figure E1.

| **Changes in gene expression** | **number of nodes** | **number of edges** | **expected number of edges** | **PPI enrichment *P*-value** |
|---|---|---|---|---|
| **Adj *P* <0.05** | | | | |
| **up and down** | 98 | 67 | 26 | 2.06E-11 |
| **up only** | 26 | 6 | 1 | 0.003 |
| **down only** | 72 | 35 | 15 | 8.52E-06 |
| **Adj *P* < 0.06** | | | | |
| **up and down** | 123 | 93 | 37 | 9.77E-15 |
| **up only** | 33 | 12 | 2 | 6.60E-06 |
| **down only** | 90 | 44 | 20 | 3.33E-06 |
| **Adj *P* < 0.08** | | | | |
| **up and down** | 156 | 133 | 55 | <1.0e-16 |
| **up only** | 46 | 18 | 5 | 8.93E-06 |
| **down only** | 110 | 55 | 26 | 3.40E-07 |

**Table E2. Summary of enriched pathways in hPAECs from pathway analysis of BMP9 regulated PPI network.**

| Changes in gene expression | enriched pathways | count in gene set | false discovery rate |
|---|---|---|---|
| **Adj P <0.05** | | | |
| **up and down** | TGFβ signaling pathway | 5 | 0.00982 |
| | cytokine-cytokine receptor interaction | 7 | 0.0367 |
| | Rap1 signaling pathway | 6 | 0.043 |
| | Hippo signaling pathway | 5 | 0.0494 |
| **up only** | Mineral absorption | 3 | 0.00927 |
| | TGFβ signaling pathway | 3 | 0.018 |
| **down only** | no enriched pathways detected | | |
| **Adj P < 0.06** | | | |
| **up and down** | TGFβ signaling pathway | 6 | 0.00201 |
| | Rap1 signaling pathway | 7 | 0.0226 |
| | cytokine-cytokine receptor interaction | 8 | 0.0226 |
| | VEGF signaling pathway | 4 | 0.0315 |
| **up only** | TGFβ signaling pathway | 4 | 0.00201 |
| | Mineral absorption | 3 | 0.00961 |
| **down only** | Rap1 signaling pathway | 6 | 0.0407 |
| | cytokine-cytokine receptor interaction | 7 | 0.0407 |
| **Adj P < 0.08** | | | |
| **up and down** | TNF signaling pathway | 8 | 0.000439 |
| | Rap1 signaling pathway | 9 | 0.00229 |
| | cytokine-cytokine receptor interaction | 10 | 0.00229 |
| | TGFβ signaling pathway | 6 | 0.00229 |
| | microRNAs in cancer | 6 | 0.0464 |
| **up only** | TGFβ signaling pathway | 4 | 0.00771 |
| | TNF signaling pathway | 4 | 0.0135 |
| | Mineral absorption | 3 | 0.0174 |
| | MicroRNAs in cancer | 4 | 0.0213 |
| **down only** | Rap1 pathway | 7 | 0.0339 |

**Table E3. Patient demographics and clinical parameters.** Enrolled subjects were characterized as systemic inflammatory response syndrome (SIRS) or sepsis by a group of blinded critical care physicians(9), or were healthy controls. A subset of randomly selected subjects from the MICU Registry with the above diagnoses was selected for analysis. Anonymized plasma samples were generated from blood collected in EDTA-containing tubes obtained from patients within 24 to 72 hours of MICU admission and stored at –80°C. Plasma was isolated from whole blood at 1500 *g* for 15 minutes at room temperature.

| Variable | **Control (n=10)** | **SIRS (n =10)** | **Sepsis (n=10)** | ***P* value** ‖ | **Statistical test** |
|---|---|---|---|---|---|
| Age (yr)* | 56 [28, 80] | 56 [28, 74] | 64 [41, 92] | 0.1364 | ANOVA with Tukey |
| Gender, n (%) | | | | 0.8663 | Chi square |
| -Female | 4 (40) | 4 (40) | 3 (30) | | |
| -Male | 6 (60) | 6 (60) | 7 (70) | | |
| Race, n (%) | | | | 0.3126 | Chi square |
| -White | 8 (80) | 6 (60) | 8 (80) | | |
| -Black | 0 (0) | 2 (20) | 0 (0) | | |
| -Hispanic | 0 (0) | 1 (10) | 1 (10) | | |
| -American Indian/Alaskan native | 0 (0) | 1 (10) | 0 (0) | | |
| -Asian/Pacific Islander | 2 (20) | 0 (0) | 1 (10) | | |
| Positive cultures, n (%) | N/A | 0 (0) | 9 (90) | <0.0001 | Chi square (0.0001 by Fisher's exact) |
| Vasopressors within 24 hr of admission, n (%) | N/A | 2 (20) | 9 (90) | 0.0017 | Chi square (0.0055 by Fisher's exact) |
| Mechanical ventilation within 24 hr of admission, n (%) | N/A | 3 (30) | 2 (20) | 0.6056 | Chi square (>0.9999 by Fisher's exact) |
| Acute Physiology and Chronic Health Evaluation (APACHE) II score* | N/A | 25 [10, 34] | 23 [19, 31] | 0.6610 | Unpaired t test |
| Sequential Organ Failure Assessment (SOFA) score* | N/A | 17 [4, 20] | 18 [13, 24] | 0.2681 | Mann Whitney |
| Comorbidities, n (%) | | | | N/A | |
| -Coronary artery disease | 0 (0) | 1 (10) | 3 (30) | | |
| -Congestive heart failure | 0 (0) | 0 (0) | 1 (10) | | |
| -Chronic obstructive pulmonary disease | 0 (0) | 1 (10) | 1 (10) | | |
| -Diabetes mellitus | 0 (0) | 5 (50) | 2 (20) | | |
| -Liver disease | 0 (0) | 3 (30) | 0 (0) | | |
| -Chronic kidney disease | 0 (0) | 2 (20) | 3 (30) | | |
| -Cancer | 1 (10) | 4 (40) | 5 (50) | | |
| -Solid tumor | 1 (10) | 2 (20) | 1 (10) | | |
| -Hematologic | 0 (0) | 2 (20) | 4 (40) | | |
| -Bone marrow transplant | 0 (0) | 1 (10) | 2 (20) | | |

| | | | | | |
|---|---|---|---|---|---|
| Creatinine† (mean+/-SD) | N/A | 2.4 ± 3.0 | 2.4 ± 1.8 | 0.3930 | Mann Whitney |
| Lactate† (mean+/-SD) | N/A | 4.3 ± 4.1‡ | 2.5 ± 1.2§ | 0.9747 | Mann Whitney |

N/A Not applicable
* Age, Acute Physiology and Chronic Health Evaluation (APACHE) II, and Sequential Organ Failure Assessment (SOFA) scores are expressed in medians [min, max 95% CI]. All other values are expressed by percentage of total of subjects (in parentheses) and total number.
† Highest value within the first 24 hours upon admission to the ICU. Creatinine units are given in mg/dL, Lactate units are given in mEq/L. Data are presented mean ± sd.
‡ $n$ = 6
§ $n$ = 8
|| $P$ values reflect overall comparison of the groups with existing data

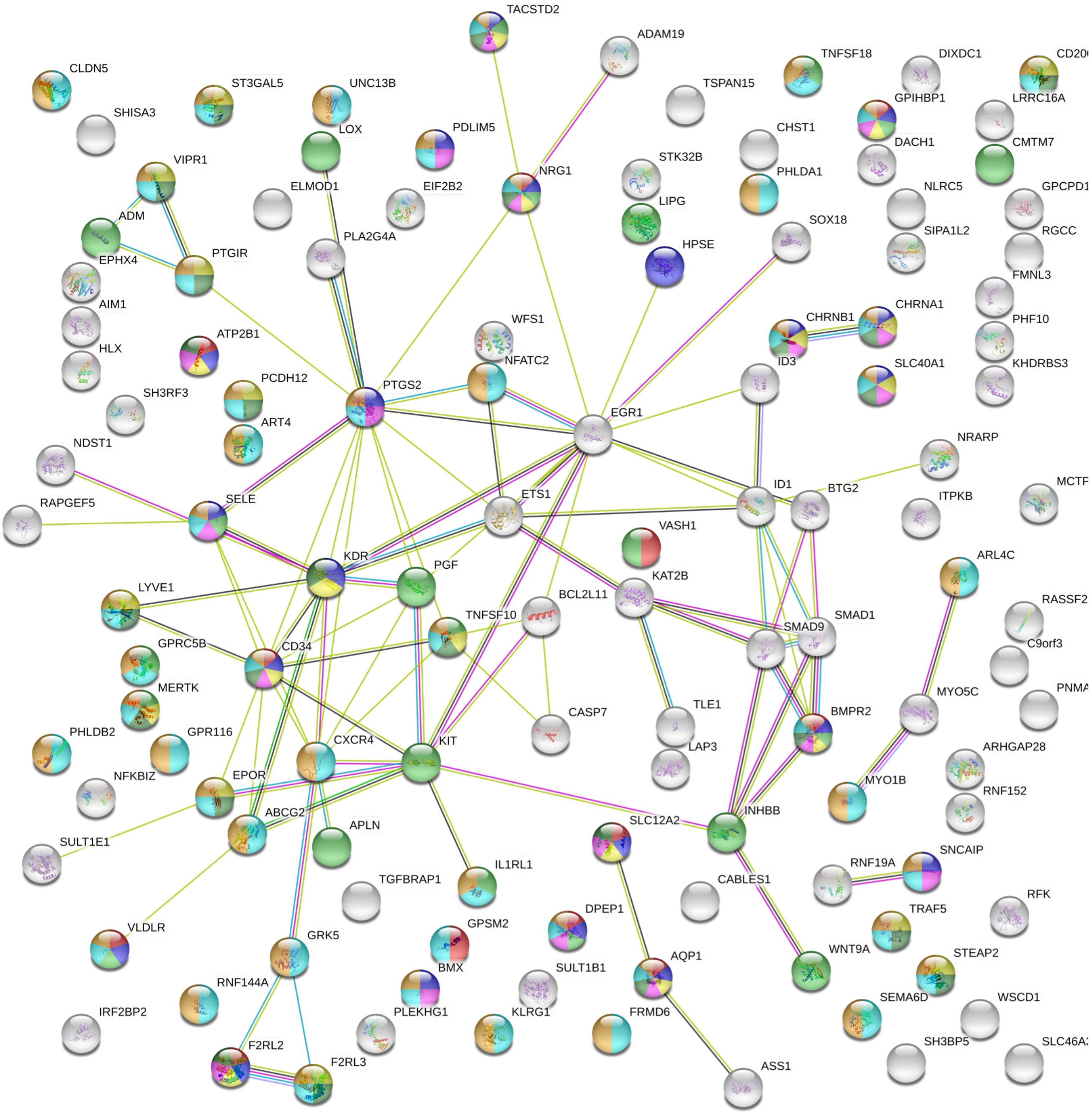

**Figure E1. BMP9 regulated genes are highly enriched on plasma membrane or extracellular space.** Protein-protein interactions network of differential regulated genes by BMP9 (using adj P-value of 0.06 cutoff, Table E1). Figure generated using STRING Sever (https://string-db.org/).

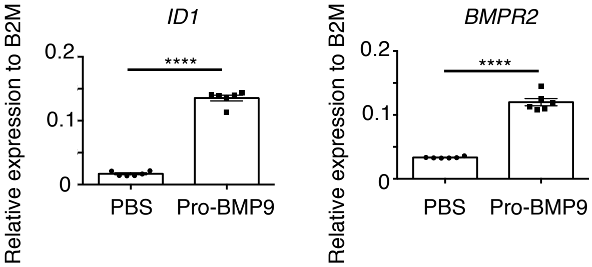

**Figure E2. BMP9 signals potently in hPMECs.** PMECs were treated with pro-BMP9 or PBS for 1.5 hours (for *ID1* gene expression) or 5 hours (for *BMPR2* gene expression) before cells were harvested for RT-qPCR analysis. N=6 independent experiments. Paired t-test, ****, P $\leq$ 0.0001.

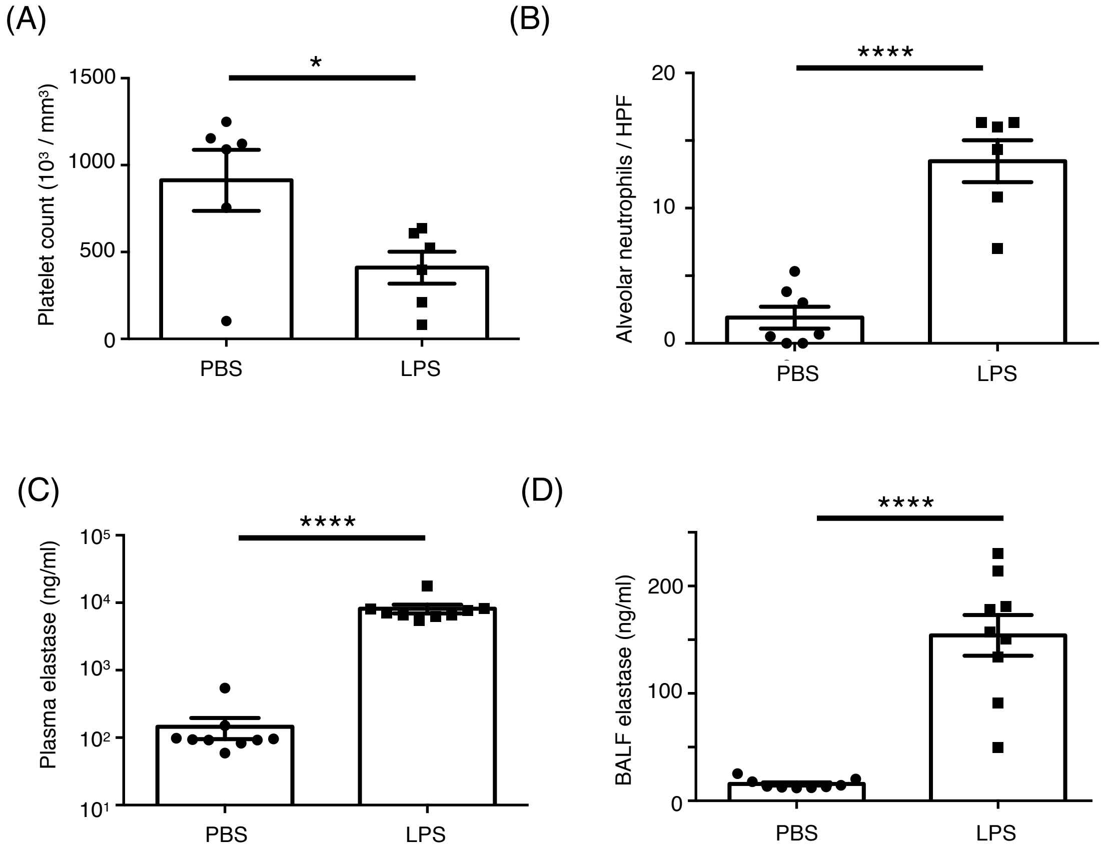

**Figure E3. Murine model of endotoxemia induced by acute intraperitoneal LPS challenge.** (A) Full blood count, showing % platelet counts were decreased in the LPS-treated animals. N=6 in each group. (B) Alveolar neutrophils were increased in the mice treated with LPS. (C&D) Elastase concentrations in plasma and BALF measured by ELISA. N=9. Data shown as means ± SEM. Unpaired T-test, *, $P < 0.05$; ****, $P \leq 0.0001$.

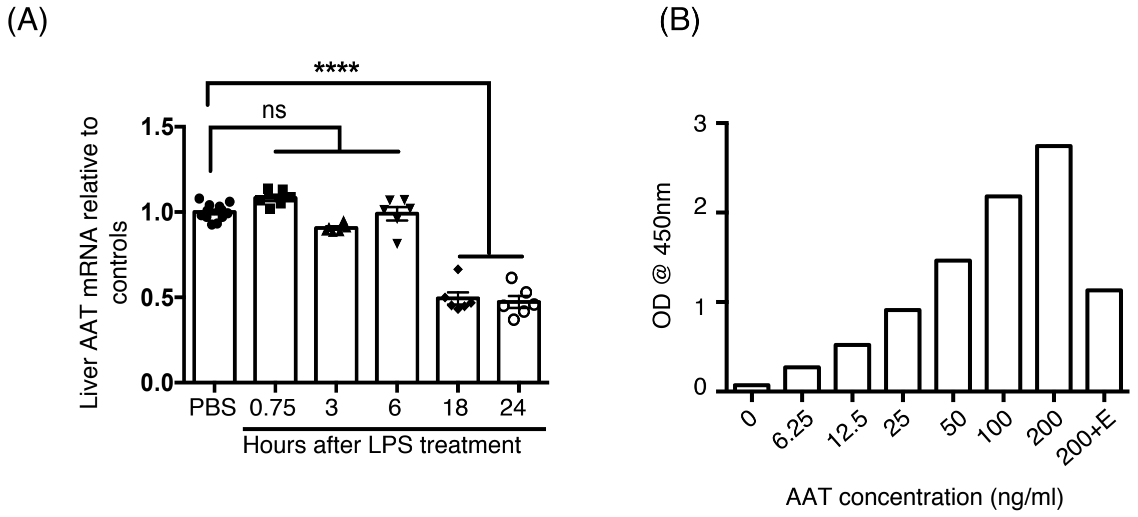

**Figure E4. The regulation of endogenous BMP9 during endotoxemia.** *(A) Changes in liver AAT mRNA relative to controls after LPS challenge. (B) Mouse AAT ELISA kit only detects native AAT.* Upon adding the elastase to the standard (200+E), the signal decreased, probably due to the large conformational change in AAT upon elastase cleavage (10).
